# The probability of chromatin to be at the nuclear lamina has no systematic effect on its transcription level in fruit fly

**DOI:** 10.1101/2022.02.17.480932

**Authors:** Alexander Y. Afanasyev, Yoonjin Kim, Igor S. Tolokh, Igor V. Sharakhov, Alexey V. Onufriev

## Abstract

Multiple studies have demonstrated a negative correlation between gene expression and the positioning of genes at the nuclear envelope (NE) lined by nuclear lamina. In this paper, we ask whether there is a causal, systematic relationship between the expression level of the groups of genes in topologically associating domains (TADs) of Drosophila nuclei and the probabilities of TADs to be found at the NE. To investigate the nature of this possible relationship, we combine a coarse-grained dynamic model of the entire Drosophila nucleus with genome-wide gene expression data; we analyze the TAD averaged transcription levels of genes against the probabilities of individual TADs to be near the NE in the control and lamins-depleted nuclei. Our findings demonstrate that, within the statistical error margin, the stochastic positioning of *Drosophila melanogaster* TADs at the NE does not, by itself, systematically affects the mean level of gene expression in these TADs, while the expected negative correlation is confirmed. The correlation is weak and disappears completely for TADs not containing lamina-associated domains (LADs) or TADs containing LADs, considered separately. Verifiable hypotheses of the underlying mechanism for the presence of correlation without causality are discussed.

## Introduction

The multi-level organization of the genome in the three-dimensional (3D) space inside the nucleus is believed to be associated with gene expression, but the exact causal connections are far from clear ***van Steensel and Furlong*** (***2019***); ***Hafner and Boettiger*** (***2019***); the importance of 3D chromatin organization for gene regulation was established in some cases ***Lupiáñez et al.*** (***2019***); ***Ibn-Salem et al.*** (***2019***); ***Spielmann et al.*** (***2019***); ***Morgan et al.*** (***2019***); ***Oudelaar et al.*** (***2019***), while in others the effect was weak or non-existent ***Ing-Simmons et al.*** (***2019***); ***Heist et al.*** (***2019***); ***Chen et al.*** (***2019***); ***Nora et al.*** (***2019***); ***Rao et al.*** (***2019***); ***Stadler et al.*** (***2019***). In particular, possible connections between gene transcription level and its position relative to the nuclear periphery have received significant attention, see, *e*.*g*., these recent reviews ***van Steensel and Belmont*** (***2019***); ***Buchwalter et al.*** (***2019***); ***Briand and Collas*** (***2019***). A degree of negative correlation has been firmly established between gene expression and its location at the periphery; however, certain aspects of the correlation are still worth investigating, especially in light of newly established properties of chromatin, such as its high mobility even for regions believed to be firmly associated with the nuclear periphery ***Kind et al. (2013***, 2015). Critically, the more difficult question of a possible causal connection, the *structure* ⟶ *function* causality, still remains open, especially if one moves beyond individual genes, and seeks to establish genome-wide, statistically significant systematic trends. A unique window of opportunity is available here to move forward by combining available experimental data with computational models, which can complement experiments with rich, genome-wide information and resolution (spatial-temporal) otherwise very difficult to obtain purely experimentally. Below is a brief review aimed to justify these claims that serve as our main motivation.

The nucleus in the eukaryotic cells is separated from the cytoplasm by the nuclear envelope (NE), the internal surface of which is lined by the nuclear lamina (NL) – a meshwork of lamins and associated proteins ***van Steensel and Belmont*** (***2019***); ***Buchwalter et al.*** (***2019***); ***Briand and Collas*** (***2019***). Lamins, as the major structural proteins of the NL, are considered to be an important determinant of nuclear architecture and gene expression ***Briand and Collas*** (***2019***). Multiple early studies have shown a correlation between positioning at the NL/NE and repression of transgenes and individual endogenous genes ***Dietzel et al.*** (***2019***); ***Misteli*** (***2019***); ***Kosak et al.*** (***2019***); ***Hewitt et al.*** (***2019***); ***Zink et al.*** (***2019***); ***Williams et al.*** (***2019***); ***Gonzalez-Sandoval and Gasser*** (***2019***). Some genes that move away from the nuclear periphery, either in lamin mutants (LM) or during tissue differentiation, have been shown to be transcriptionally upregulated ***Williams et al.*** (***2019***); ***Malhas et al.*** (***2019***). However, expression of other genes appears unaffected by their proximity to the nuclear periphery ***Nielsen et al.*** (***2019***); ***Zhou et al.*** (***2019***); ***Hewitt et al.*** (***2019***). The consistency of gene contacts with NL was shown to be negatively correlated with the gene activity (and positively with heterochromatic histone modification H3K9me3) in single cells of in the human myeloid leukemia cell line KBM7 ***Kind et al.*** (***2019***). A weak genome-wide negative correlation has been found between expression and NL-contacts of genomic regions that infrequently associate with the nuclear periphery ***Rooijers et al.*** (***2019***). The small effect size between the association of genome–NL contacts with transcription could be the result of the limited time resolution of these experiments (12 h) ***Rooijers et al.*** (***2019***). Genomic regions associated with NL can have highly expressed genes in developing mouse brains, suggesting that NL binding by itself does not repress transcription ***Ahanger et al.*** (***2019***). Similarly, genome-wide disassociation of genomic regions from NL in *Caenorhabditis elegans* caused upregulation of only a single gene ***Gonzalez-Sandoval et al.*** (***2019***).

Several studies directly tested functional consequences of chromatin-lamina association by tethering individual genes to the nuclear periphery in human cells using the lacO/LacI system ***Finlan et al.*** (***2019***); ***Kumaran and Spector*** (***2019***); ***Reddy et al.*** (***2019***). Since binding of nucleoplasmic LacI molecules to lacO sites in the reporter construct did not, by itself, impair transcription ***Finlan et al.*** (***2019***); ***Reddy et al.*** (***2019***), these studies provided direct evidence for a causative role of the nuclear periphery in altering gene expression of many genes. For example, 51 endogenous genes located around a site of induced NE-attachment have been repressed, suggesting that an inactive chromosomal domain has been generated upon tethering to the lamina ***Reddy et al.*** (***2019***). Notably, the transcriptional repression caused by the tethering of a gene to the NE was accompanied by histone H4 hypo-acetylation, implying the importance of epigenetic modifications in this process, not necessarily the geometric proximity to the NE *per se*. At the same time, some tethered genes were not repressed ***Finlan et al.*** (***2019***); ***Reddy et al.*** (***2019***). Moreover, other experiments have demonstrated that lamina-targeted genetic loci can still be activated and transcribed at the lamina ***Kumaran and Spector*** (***2019***). The tethering studies suggest that, for some genes, the NL represents a repressive compartment in the cell nucleus, but the repression can be overcome by other genes. Overall, these works discovered important features of individual genes and groups of genes, but they did not make genome-wide conclusions.

More recently, chromatin regions called lamina-associated domains (LADs), were systematically identified using a DamID approach in cell nuclei of humans and fruit flies ***Pickersgill et al.*** (***2019***); ***Guelen et al.*** (***2019***); ***van Bemmel et al.*** (***2019***); ***Pindyurin et al.*** (***2019***). Attachment of LADs to the NL is stochastic: only about 25-30% of all LADs in the fruit fly are located at the nuclear periphery at any given moment in any given cell ***Pickersgill et al.*** (***2019***); ***van Bemmel et al.*** (***2019***); ***Ulianov et al.*** (***2019***). Similar to other chromosome loci in the interphase nuclei ***Marshall et al.*** (***2019***); ***Csink and Henikoff*** (***2019***); ***Chubb et al.*** (***2019***); ***Spector*** (***2019***); ***Lanctôt et al.*** (***2019***), the fruit fly LADs are highly mobile within each nucleus. Despite having a relatively strong affinity to the NL ***Tolokh et al.*** (***2019***), most fruit fly LADs come in contact with the NL and move away from it multiple times during the interphase. Therefore, tethering experiments in which a locus is permanently anchored to the NL, or DamID studies with a limited temporal resolution of NL-contacts during the interphase, may not represent the highly mobile nature of chromatin in fruit flies. Forced expression of several genes located mostly at the nuclear periphery can cause their detachment from the NL ***Therizols et al.*** (***2019***). Genome-wide gene activation inside LADs typically causes detachment of LADs from the NL, but it usually does not involve more than 50 kb of DNA flanking the activated gene, and the NL detachment is dependent on transcription elongation ***Brueckner et al.*** (***2019***).

An experiment that can probe, directly, whether a statistically meaningful change in the necessarily stochastic position of a gene relative to the NE causes a systematic, statistically meaningful change in the gene expression (the *structure* ⟶ *function* causality) would be most appropriate, but to the best of our knowledge, no such studies have been performed on a genome-wide scale. For example, a recent pioneering genome-wide analysis of gene transcriptions in control vs. Lamin-knockdown (Lam-KD) fruit fly nuclei ***Ulianov et al.*** (***2019***) demonstrated that, upon Lam-KD, a number of genes in LADs increased their transcription levels by up to a factor of 4, while for another subset of genes, a decrease by up to a factor of 2 was observed. However, changes in the cumulative spatial distributions of the chromatin regions upon Lam-KD which were revealed for only three genomic regions in LAD-containing TADs, which speaks for the difficulty of performing such experiments for every gene.

The difficulties in performing this kind of analysis purely experimentally motivate combining experiment with an appropriate computer modeling ***Liang and Perez-Rathke*** (***2019***); ***Tolokh et al.*** (***2019***); ***Contessoto et al.*** (***2019***) to make progress. Here, an experiment can provide gene expression data, while computer modeling can trace the movement of the relevant chromatin units with the desired spatial and temporal resolution on a genome-wide scale to detect systematic trends if any.

Due to the highly stochastic nature of transcription at the level of individual genes, relatively large units of chromatin structure—beyond an individual gene—are more appropriate for the goal of exploring systematic trends in gene expression by averaging over many genes that are close to each other in 3D space. Our specific choice is introduced below.

Application of the chromosome conformation capture technique with high-throughput sequencing (Hi-C) to study chromatin organization identified topologically associating domains (TADs) in various organisms of high eukaryotes ***Lieberman-Aiden et al.*** (***2019***); ***Dixon et al.*** (***2019***); ***Nora et al.*** (***2019***); ***Sexton et al.*** (***2019***); ***Ulianov et al.*** (***2019***). TADs may contain multiple genes, range in size from tens of kilobases (kb) in fruit flies to several megabases (Mb) in mammals, and represent structural and functional units of three-dimensional (3D) chromatin organization ***Sexton and Cavalli*** (***2019***); ***Gorkin et al.*** (***2019***); ***Rowley and Corces*** (***2019***); ***Yasuhara and Zou*** (***2019***). Since the boundaries between TADs are conserved among cell types and sometimes across species ***Dixon et al.*** (***2019***); ***Phillips-Cremins et al.*** (***2019***); ***Rao et al.*** (***2019***); ***Dixon et al.*** (***2019***); ***Acemel et al.*** (***2019***); ***Ulianov et al.*** (***2019***); ***Renschler et al.*** (***2019***); ***Torosin et al.*** (***2019***); ***Liao et al.*** (***2019***), a TAD can be regarded as a natural unit of genome partitioning for exploring systematic structure-function relationships in chromatin. Certain TADs overlap with LADs in cell nuclei of humans and fruit flies ***Guelen et al.*** (***2019***); ***van Bemmel et al.*** (***2019***); ***Pindyurin et al.*** (***2019***); ***Sexton et al.*** (***2019***); ***Nora et al.*** (***2019***); ***Dixon et al.*** (***2019***).

In this work, we combine published ***Ulianov et al.*** (***2019***) genome-wide gene transcription data with a novel modeling approach to ask whether there is a causal, systematic relationship between the transcription level of the groups of genes in TADs of *Drosophila* nuclei and the probability of TADs to be found at the NE, taking into full account the highly mobile nature of the TADs. For this purpose, we employ a recently developed model of the fruit fly interphase chromatin ***Tolokh et al.*** (***2019***) that describes the dynamics and spatial organization of the entire diploid set of female chromosomes at TAD resolution, and their interactions with the NE. The model, which accounts for different epigenetic classes of TADs, TAD-TAD contact probabilities (Hi-C map), and the known distribution of LADs along the genome, reproduces experimentally observed chromatin density profiles for both control and lamins-depleted nuclei and can faithfully predict the probabilities of individual TADs to be in the layer adjacent to the NE. To investigate the genome-wide relationship between the positioning of TADs at the NE and gene transcription at TAD resolution, a robust TAD-averaged metric of transcription activity is introduced.

## Results

### TAD transcription levels and the probability of a TAD to be near the NE: correlation

Here, we investigate whether average gene transcription levels in TADs, that are relatively free to move within the fruit fly interphase nuclei ***Marshall et al.*** (***2019***); ***Vazquez et al.*** (***2019***); ***Lanctôt et al.*** (***2019***); ***Tolokh et al.*** (***2019***), correlate with TADs average geometric proximity to the NE. While we do expect such a correlation, in general, ***Shachar and Misteli*** (***2019***), we aim to investigate it in detail; for example, it may exist only for a certain subgroup of TADs. As the next step, we would like to test whether the positioning of TADs at the NE *per se* causes the change in transcription.

To this end, we need a probabilistic metric for the TAD-NE proximity, because the same TAD can move in and out of proximity to the NE multiple times during the interphase ***Tolokh et al.*** (***2019***). It is reasonable to assume that, if proximity to the NE suppresses transcription, the degree of suppression of gene transcription activities in a TAD should be related to the fraction of time the TAD spends in the proximity to (*i*.*e*., in contact with) the NE. This fraction can be expressed as the probability of a TAD to be in a rather narrow contact layer near the NE, that is when its center is within 0.2 *μ*m (average bead diameter in the model ***Tolokh et al.*** (***2019***)) from the NE, see Materials and Methods and Figure 1 of Appendix 1. We consider this probability (frequency) of a TAD to be in this layer near the NE, that is, to be essentially in contact with the NE, as a suitable measure of the TAD’s *stochastic* proximity to the NE.

**Figure 1.**
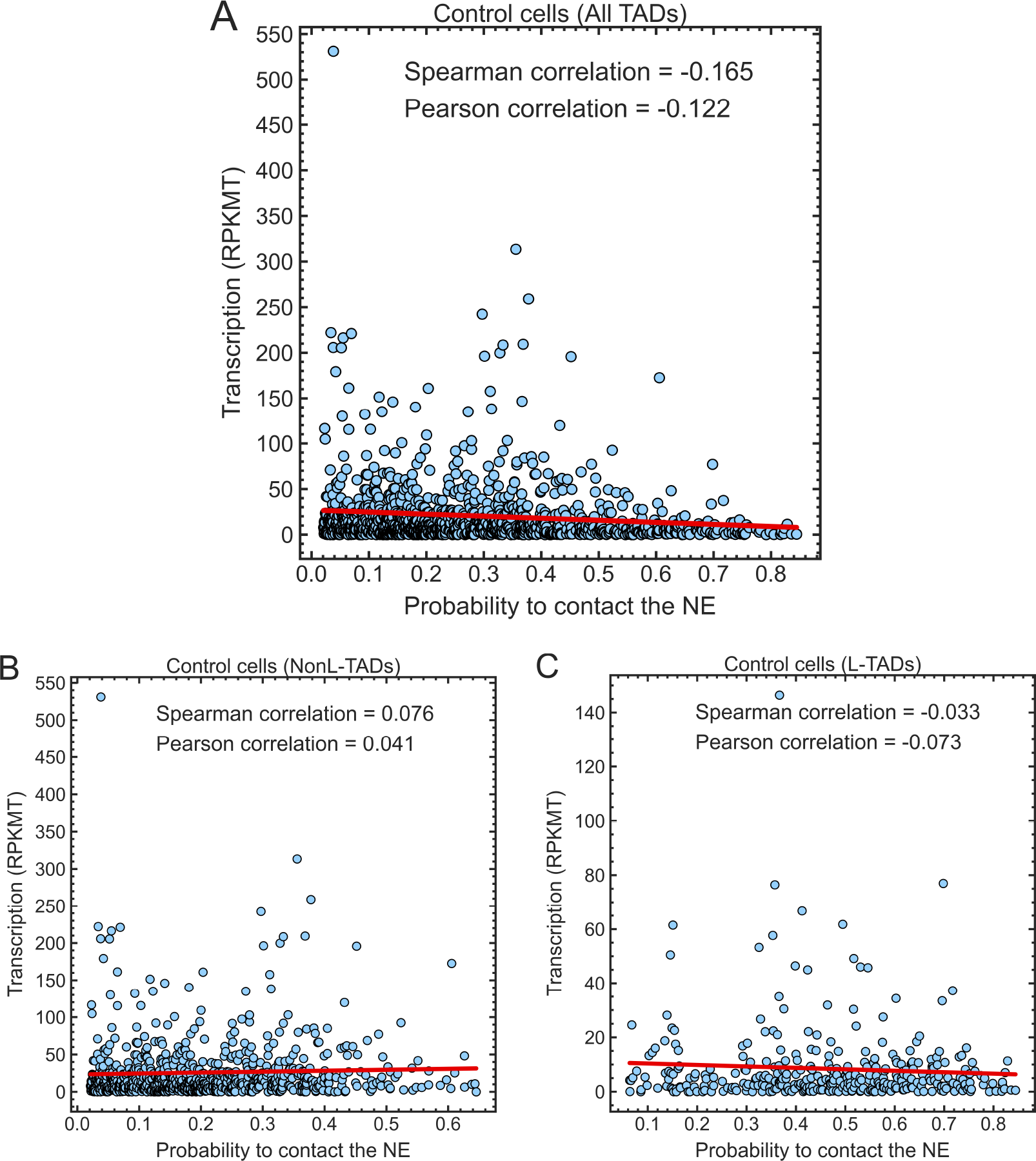
Transcription levels, in RPKMT, of genes in each TAD vs. the probability of a TAD to be in contact with the NE in the control cells. **(A)** A weak negative correlation is seen when all TADs are considered. **(B and C)** The scatter plots show essentially no correlation for NonL-TADs or L-TADs, considered separately. The Spearman and Pearson correlation coefficients, and linear regression lines (red) are shown. Plots are generated with the *lmplot* function in Seaborn.

In this work, we introduce and employ a normalized metric for gene transcription levels at TAD resolution, RPKMT. This metric extends the commonly used single gene transcription level metric, RPKM (reads per kilobase of transcript per million reads mapped), to TAD resolution reflecting a TAD averaged transcription activity, see Materials and Methods. To make sure that our main conclusions are robust to details of the transcription metric definition, we also consider another metric of transcription levels in TADs, RPKMTL (number of reads mapped to all genes in a TAD per kilobase of TAD length per million reads mapped to all TADs), see Appendix 1. In this metric, a TAD length instead of a sum of gene lengths in a TAD is used.

Figure 1A shows that there is a weak negative correlation between the transcription activity in TADs and their probability to be in contact with the NE in the control (*i*.*e*., with the intact lamina) *Drosophila* cells. The absence of a pronounced negative correlation may be unexpected, as the transcription of genes in LADs and, therefore, in TADs containing LADs (L-TADs), which interact with the NL and constitute a significant (about 30%) fraction of TADs in *Drosophila* ***van Bemmel et al.*** (***2019***); ***Sexton et al.*** (***2019***), is well known to be repressed ***Reddy et al.*** (***2019***); ***Kim et al.*** (***2019***); ***Briand and Collas*** (***2019***). Nuclear periphery is generally assumed to be an inactive compartment ***Lanctôt et al.*** (***2019***).

To gain further insight into the weak correlation seen in the control cells nuclei (Figure 1A), we separate the TADs into two major groups: TADs not containing LADs (NonL-TADs) and TADs containing LADs (L-TADs), see Figure 1B and C. We see essentially no correlation between the transcription levels (RPKMT) in L-TADs/NonL-TADs and the probability of L-TADs/NonL-TADs being in contact with the NE in control cells. Much lower average gene transcription levels in L-TADs relative to NonL-TADs (compare the levels of linear regression lines in Figures 1B and C) and the absence of NonL-TADs with the probabilities to be in contact with the NE greater than 0.65 can explain the weak negative correlation between the transcription levels and the “contact” probabilities for all TADs seen in Figure 1A.

Given the already weak correlation between the transcription levels of TADs (in RPKMT) and the probabilities of TADs to be in contact with the NE, seen in the control nuclei, it is not surprising that no correlation between the transcription activities in TADs and their contact probabilities is seen in the lamins-depleted nuclei, where fewer TADs are expected to be in contact with the NE relative to the control cells, Figure 2. In what follows, we investigate whether a weak signal is still there, “drowned out” by the inherently noisy data.

**Figure 2.**
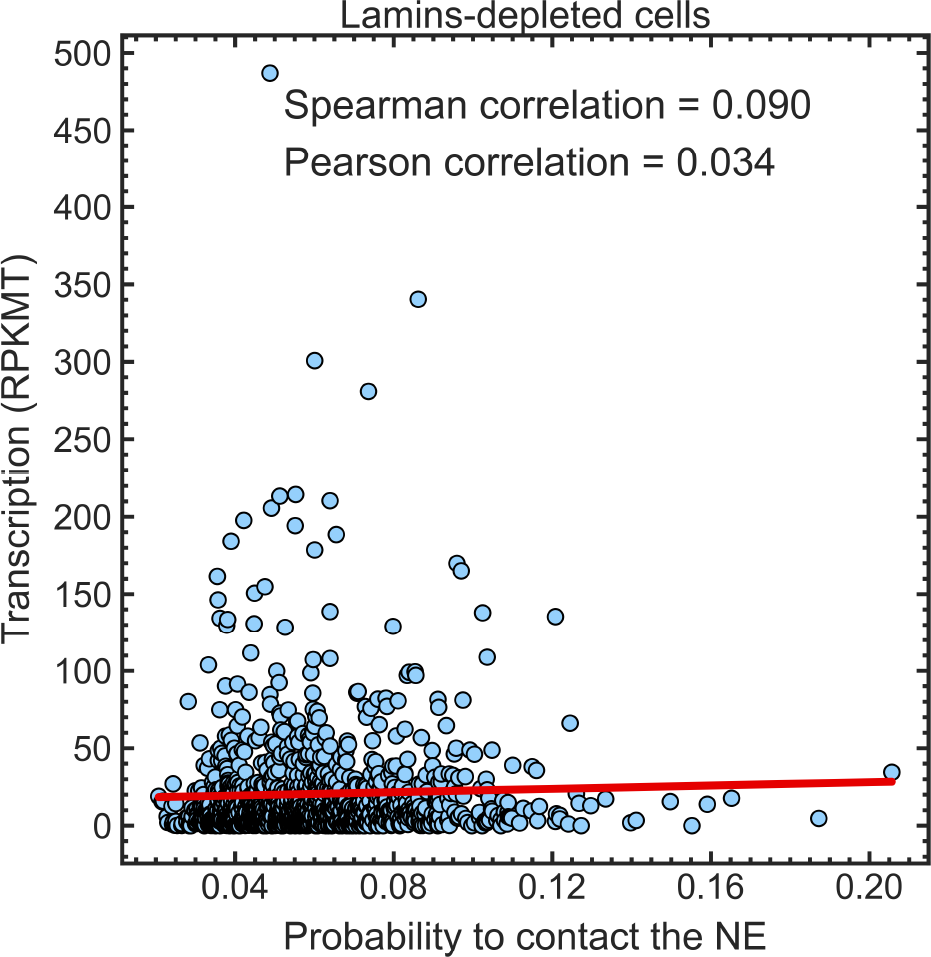
Essentially no correlation is seen between the transcription levels in all TADs (in RPKMT) in the lamins-depleted cells and the probability of TADs being in contact with the NE. The Spearman and Pearson correlation coefficients, and the linear regression lines (red) are shown. Plots are generated with the *lmplot* function in Seaborn.

To discern meaningful, systematic, genome-wide trends behind the large natural variation of transcription activity between individual TADs, as well as inevitable errors in the experimentally reported gene transcription levels, we propose averaging transcription levels even beyond individual TADs, which will help us to focus on systematic trends. We also argue for comparisons of mean values directly, rather than of means of ratios ***Ulianov et al.*** (***2019***), as the latter can contain unintended biases towards higher values due to inevitable uncertainties, see the discussion next to Table 1 of Appendix 1.

To this end, we have binned the TADs (separately all TADs, L-TADs and NonL-TADs) based on the probability of a TAD in control cells to be in contact with the NE (see Materials and Methods). Each of the six bins in Figure 3A (B,C) contains an approximately equal number of TADs (see Materials and Methods). The heights of the bars indicate the average transcription level of TADs (in RPKMT) in each bin.

**Figure 3.**
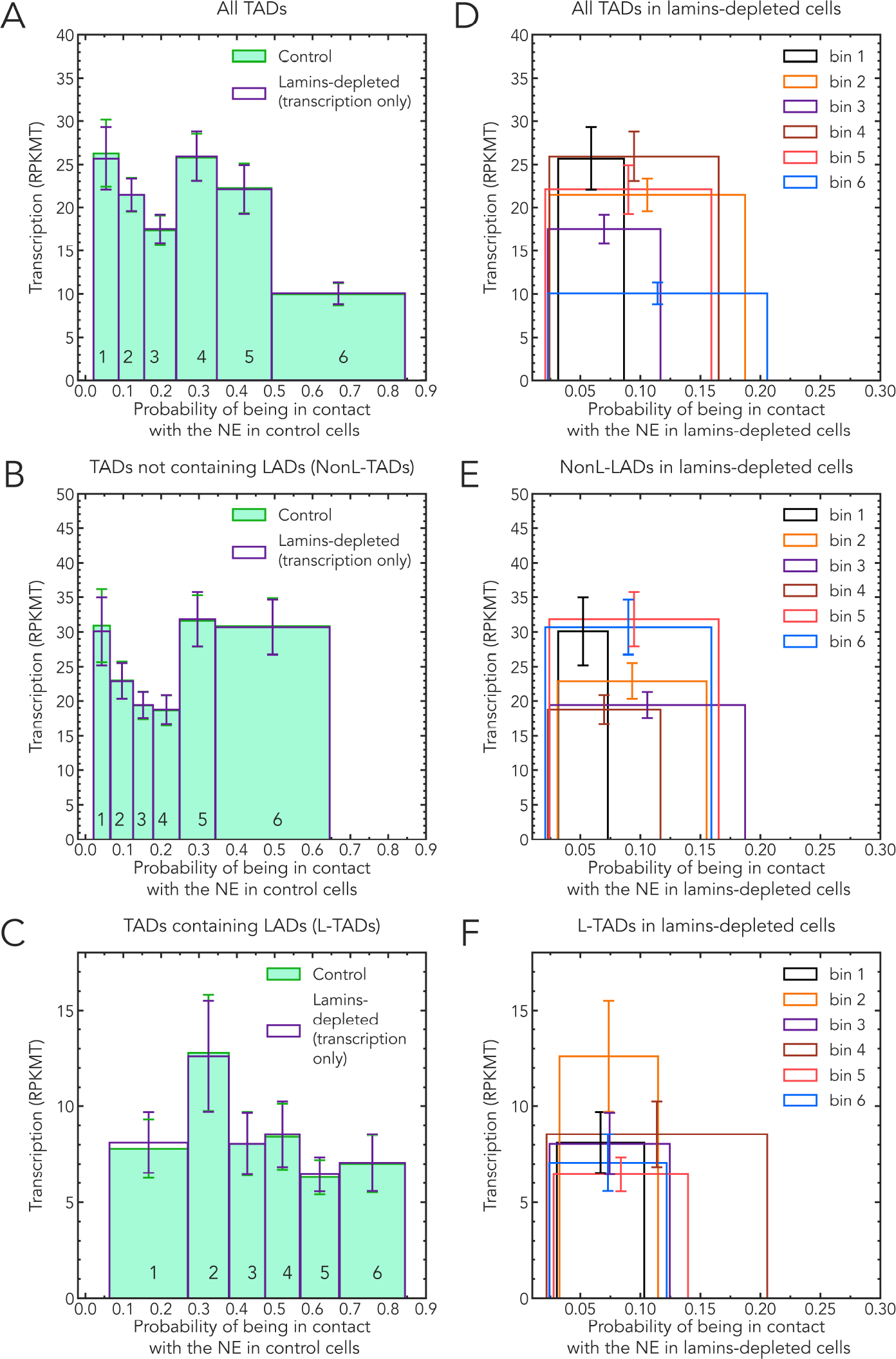
Dependencies of bin averaged TAD transcription levels (RPKMT) on the average probability of the TADs in the bin to be in contact with the NE. The binning of TADs is based on TAD-NE contact probabilities in control cells for each set (selection) of TADs. **Solid bars: control cells. Empty bars: lamins-depleted cells**. The same set of TADs per bin is used in the control and lamins-depleted cells. Error bars are s.e.m. (standard error of the mean). **Left panels: (A)** all TADs; **(B)** TADs not containing LADs (NonL-TADs); and **(C)** TADs containing LADs (L-TADs). In the left panels only, the positions of the empty bins (lamins-depleted cells) along the x-axis are deliberately kept unchanged to facilitate visual comparison with the heights of the corresponding control bins. **Right panels: (D)** all TADs; **(E)** TADs not containing LADs (NonL-TADs); and **(F)** TADs containing LADs (L-TADs). A clear shift of the average TAD positions away from the NE is evident.

Considering all TADs (i.e., both NonL-TADs and L-TADs, Figure 3A), the TADs in bin #6 – TADs that are most likely to be in contact with the NE – do show a marked (2x) decrease in transcription levels compared to TADs in other bins with a lower probability to be in contact with the NE. This trend, however, does not continue for TADs in bin #5, with the next highest level of contact probabilities. The average transcription level in bin #5 is roughly the same as those of bin #1 and #2 that contain TADs with the very low (less than 0.15) contact probabilities.

As we suggested earlier, analyzing data in Figure 1, the abrupt drop in the transcription seen in bin #6 is due to a very high fraction of L-TADs in this bin (see Figure 4). These LAD-containing TADs have 2 to 4 times lower average TAD transcription levels compared to NonL-TADs (see Figures 3B and 3C, and Ref. ***Ulianov et al.*** (***2019***)). Although a small fraction of genes located in LADs can actively express (about 10%), transcription of the majority of genes inside LADs is repressed to a very low level ***Pickersgill et al.*** (***2019***); ***Guelen et al.*** (***2019***); ***van Bemmel et al.*** (***2019***); ***Zheng et al.*** (***2019***); ***Wu and Yao*** (***2019***); ***Ulianov et al.*** (***2019***), leading to relatively low levels of transcription in L-TADs. No correlation of the average transcription levels with the likelihood to be in contact with the NE is seen for NonL-TADs in Figure 3B. The drops in the transcription levels seen in bins #3 and #4 (Figure 3B) are tangential to the main focus of this work, and so we do not pursue their origins here; a relevant hypothesis is outlined in Appendix 1, Figure 7.

**Figure 4.**
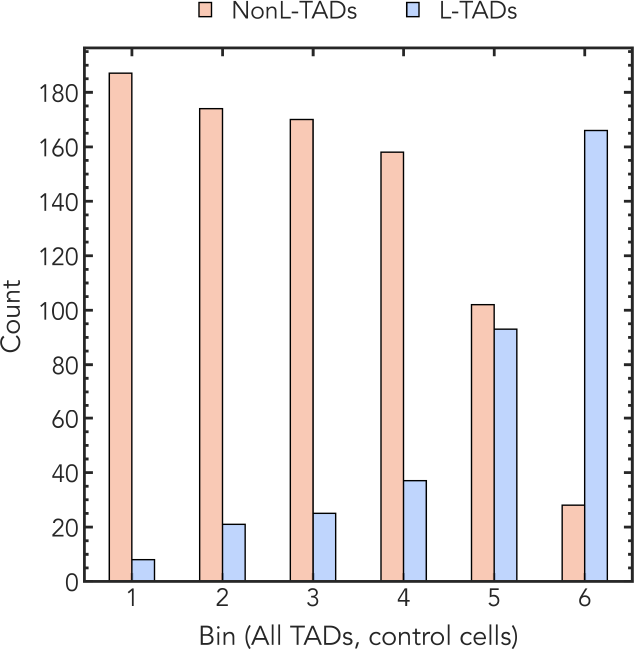
Comparison of the number of L-TADs and NonL-TADs in each bin of control cells for All TADs. The number of L-TADs in each bin increases with the proximity to the nuclear envelope.

In summary, our first conclusion is that the weak negative correlation between the transcription level of all TADs and their stochastic proximity to the NE in the control cells is due to an exceptionally large fraction of L-TADs among TADs with the highest probabilities (greater than 0.5) to be in contact with the NE and much lower (3-4 times) average transcription levels in L-TADs compared to NonL-TADs.

### TAD transcription levels vs. the probability of a TAD to be near the NE: no causation

A natural question arises whether the weak negative correlation between the transcription activity of TADs and their stochastic proximity to the NE in fruit fly nuclei is causal. In other words, does the increased probability to find a TAD in contact with the NE, by itself, cause the transcription level to decrease?

To address this question, we have considered the transcription patterns in lamins-depleted cells. First, note that the lamins knockdown or a mutation causes a drastic spatial re-arrangement of chromatin within the nucleus ***Ulianov et al.*** (***2019***); ***Bondarenko and Sharakhov*** (***2019***); ***Tolokh et al.*** (***2019***) – most of the TADs move away from the NE, see Figures 3D,E,F, as well Figure 2 of Appendix 1. The aforementioned LamA25 mutation removes the CaaX motif from Lamin B and disrupts its attachment to the nuclear membrane, thus, depleting Lamin B from the nuclear periphery ***Bondarenko and Sharakhov*** (***2019***); ***Patterson et al.*** (***2019***). In fact, the probability to find most of the TADs in contact with the NE in lamins-depleted nuclei is less than 0.16, with no bins extending beyond 0.21, Figures 3D,E,F. See also Figure 1 of Appendix 1 for the stark contrast between the control and lamins-depleted cells in this respect. If being in contact with the NE with an appreciable probability caused a *systematic* drop in the transcription of TADs containing LADs, the average transcriptional activity of these TADs would be expected to recover almost fully in lamins-depleted nuclei. Yet, no change in average transcription levels is seen in each of the *same* groups of TADs (bins), Figure 3, open bars (lamins-depleted) vs. solid bars (control). Note that we deliberately kept exactly the same sets of TADs in each lamins-depleted bins as in the control bins.

Thus, our second and main conclusion is that stochastic proximity to the NE of TADs, which are highly mobile, does not, by itself, cause a noticeable *systematic, genome-wide* change in transcription at TAD resolution. The key conclusions mentioned above are robust with respect to the metric of transcription activity used, see Figure 3,4 of Appendix 1, where an alternative metric was employed.

## Discussion

This work has combined simulation and experiment to arrive at the following main result: the probability of a TAD to be found in contact with the NE, by itself, has no systematic, causal effect on its transcription level in *Drosophila*. We stress several key aspects of this statement. First, it applies to a systematic trend only, averaged across multiple TADs, and not necessarily to experiments in which individual TADs are perturbed, e.g. permanently tethered to the NE. As shown in ***Ulianov et al.*** (***2019***), it is entirely possible for transcription levels of a subset of individual, weakly expressing TADs to increase in lamins-depleted nuclei, where their probability to be in contact with the NE is expected to decrease relative to the control nuclei. There is no contradiction here with our main claim: even if more TADs become transcriptionally up-regulated then down-regulated upon lamins depletion, which was indeed seen in Ref. ***Ulianov et al.*** (***2019***), the net effect averaged over large groups of TADs can still be zero. And this is what we are finding, within the statistical margin of error. In addition, lamins knockdown used in ***Ulianov et al.*** (***2019***) can affect gene expression in more than one way, including an increase of the acetylation level at histone H3 and a decrease of chromatin compaction in LADs.

Second, we stress the highly mobile nature of TADs ***Tolokh et al.*** (***2019***), including L-TADs, reflected in our deliberate word choice of “probability to be in contact with the NE” *vs*. the more common “proximity to the NE”. We have also used “stochastic proximity” to emphasize the same point. L-TADs, even those that are likely to be found in contact with the NE, move away from the NE multiple times during the interphase ***Tolokh et al.*** (***2019***). Thus, the condition of a stable attachment to the NE, *e*.*g*., *via* a tether, is not reproduced in live *Drosophila* nuclei, which removes the perceived contradiction with the pioneering tethering studies that demonstrated down-regulation of certain loci upon induced contact with the NE ***Finlan et al.*** (***2019***); ***Kumaran and Spector*** (***2019***); ***Reddy et al.*** (***2019***). Indeed, targeting of lamin-associated loci does not take place during interphase when loci can only form transient contacts with the lamina ***Kumaran and Spector*** (***2019***). Stable tethering of a locus to the nuclear lamina requires passage through mitosis. Similarly, treatment of *Drosophila* S2 cells with dsRNA against lamin Dm0 was performed over four days ***Ulianov et al.*** (***2019***) allowing passages through mitosis. It is likely that cell division is necessary for the re-setting of an epigenetic state of the chromatin at the nuclear periphery. Thus, epigenetic marks and transcription levels of TADs are unlikely to change during the same interphase, despite the fact that TADs frequently change their positions with respect to the lamina.

Different mechanisms for LAD positioning and repression are not expected to be mutually exclusive. For example, a study performed a detailed analysis of gene repression mechanisms in LADs using human K562 cells ***Leemans et al.*** (***2019***). By systematically moving promoters from their native LAD location to a more neutral chromatin environment and by transplanting them to a wide range of chromatin contexts inside LADs, the study has demonstrated that the variation in the expression level can be due to the interplay between the promoter sequence and local chromatin features in the LADs ***Leemans et al.*** (***2019***). Chromatin features in the LADs can be partially responsible for interaction with the lamina since lowering H3K9me2/3 levels decreases LAD-contacts in human cells ***Kind et al. (2013***, 2015). The lamina can attract loci of inactive chromatin, thus, contributing to the assembly of repressive peripheral chromatin domains. The NE proteins can also attract chromatin to the lamina and participate in the repression of transcription. For example, histone deacetylase HDAC3, which is associated with the NE transmembrane (NET) protein Lap2beta and the DNA-binding protein cKrox, was shown to attract LADs to the nuclear periphery in mice cells ***Zullo et al.*** (***2019***); ***Poleshko et al.*** (***2019***) and Drosophila S2 cells ***Milon et al.*** (***2019***). Inhibition of histone deacetylation with trichostatin A (TSA) in *Drosophila* cells causes loss of Lamin binding to chromatin ***Pickersgill et al.*** (***2019***). Thus, nuclear lamina-associated HDACs could contribute to the repressive environment of the nuclear periphery by downregulating gene expression ***Somech et al.*** (***2019***); ***Demmerle et al.*** (***2019***) and assembling the peripheral chromatin. Moreover, lamins themselves can affect both the position and transcription of chromatin. A study has shown that artificial anchoring of lamins to promoters of transfected reporter plasmids can lead to reduced transcription ***Lee et al.*** (***2019***). A conditional and temporal up-regulation of lamin C in *Drosophila* caused a shift in chromatin distribution from peripheral to central, which was associated with reduced levels of the active chromatin mark H3K9ac ***Amiad-Pavlov et al.*** (***2019***). Depletion of lamin Dm0 resulted in the moderate upregulation of the generally very weak background transcription in LADs but not in the inter-LADs in *Drosophila* S2 cells ***Ulianov et al.*** (***2019***). Also, lamin B1 depletion decreased heterochromatin marker H3K27me3 by 80 percent in human cells ***Stephens et al.*** (***2019***). While confirming the well-known repressive role of the NE, we make a conclusion here that the expected negative correlation between the TAD transcription level and its stochastic proximity to the NE disappears completely for lamina-associated domains (LADs) or non-LADs, considered separately. Finally, we believe that our general conclusions can be extended to mammals, with caution. While TADs and LADs exist in both cases, unlike mammals, TAD boundaries in fruit flies are not associated with CTCF ***Ulianov et al.*** (***2019***); ***Cubeñas-Potts et al.*** (***2019***); ***Eagen et al.*** (***2019***) and LAD-NL binding in *Drosophila* is not Lamin B Receptor dependent ***Ulianov et al.*** (***2019***). Nevertheless, functions of TADs, LADs and chromatin compartments are similar among different organisms ***Manzo et al.*** (***2019***); ***Shevelyov and Ulianov*** (***2019***); ***Solovei et al.*** (***2019***); ***Acemel and Lupiáñez*** (***2019***); ***Lukyanchikova et al.*** (***2019***).

Our study has several limitations. The conclusions rely on predictions of a computational model: while the systematic, and even a few individual trends in the predicted TAD positioning have been verified against experiment, deviations from reality may still occur on the level of individual TADs. Thus, we stress the systematic nature of the main conclusion, likely correct in the average sense. A related limitation is that the minimal structural unit of chromatin employed in this work is a TAD, which is about 100 kb for *Drosophila*. We do not make any claims about what might be happening at finer resolutions, including promoter-enhancer interactions within TADs. We stress that the resolution limitation does not invalidate our main conclusions based on averages. For example, it is known that individual active genes within a LAD can locally detach from the lamina. But still, genes belonging to a LAD that is stochastically closer to the NE are, on average, closer to the NE than genes in a LAD that is stochastically further away from the NE. Therefore, conclusions that rely on averages over these genes remain valid. Another possible limitation is that the cell types used to define the epigenetic classes employed by the computational model and to generate the transcription profiles (RNA-seq) were taken from related, but not exactly the same cell cultures. The concern is mitigated by the fact that both cell types have an embryonic origin. Moreover, we have confirmed the consistency of the transcription profiles within the four epigenetic classes of TADs between S2 embryonic cell line data ***Ulianov et al.*** (***2019***) used in our study, Figure 5A, and 16-18 hr embryos data ***Sexton et al.*** (***2019***) presented in Figure 5B. The data from this latter work (***Sexton et al.*** (***2019***)) were used to develop parameters of the computer model ***Tolokh et al.*** (***2019***) from which we compute the TAD-NE contact probabilities.

**Figure 5.**
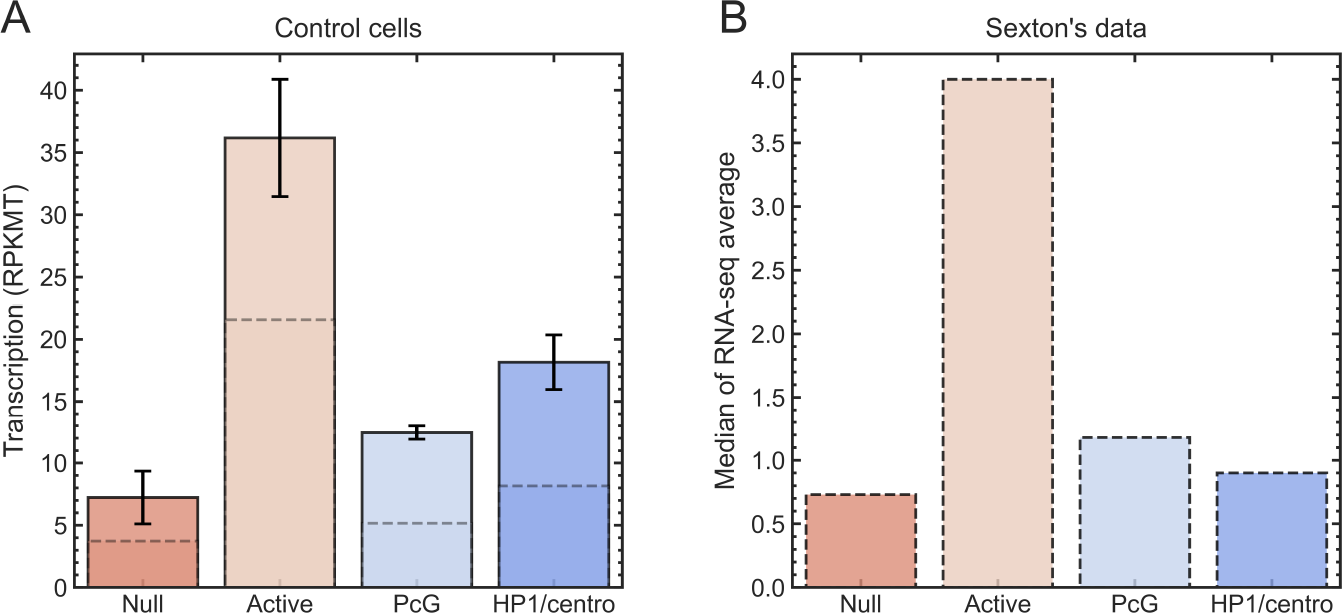
The proposed metric for transcription activity in TADs, RPKMT, is consistent with the transcription activities in different epigenetic classes of TADs ***Sexton et al.*** (***2019***). **Panel A:** Mean (solid boxes) and median (dashed lines) transcription levels (in RPKMT) for Null (n=492), Active TADs (n=494), PcG (n=131) and HP1/centromeric (n=52) epigenetic TAD classes. Error bars are standard errors of the mean. Both mean and median RPKMT transcription levels in Active TADs are at least 2 times greater than those in the three repressive epigenetic TAD classes (Null, PcG and HP1/centromeric). These RPKMT transcription levels demonstrate consistency with the gene transcription levels (in medians of RNA-seq averages) within each epigenetic class of TADs shown in Figure 3C of Ref. ***Sexton et al.*** (***2019***), reproduced in **panel B**.

The fourth limitation is that the lamin knockdown may not only cause global chromatin relocation, but also epigenetic perturbation of TADs. This concern is mitigated by the fact that the level of histone H3 acetylation is elevated only in LADs, but not in the LAD-free regions upon Lamin knockdown when compared to control cells ***Ulianov et al.*** (***2019***). Thus, the epigenetic profiles of the majority of TADs are expected to be unchanged upon the Lamin knockdown.

### Ideas and Speculative Predictions

The main implication of this work is that, with some relatively rare exceptions, the role of the NE does not include significant *systematic* regulation of transcription states of genes in TADs. Instead, the lamina may help to “lock in” the repression state of L-TADs by more than one mechanism. First, the nuclear lamina may facilitate better separation of active and inactive chromatin by sequestering inactive TADs (Null, PcG, and HP1/centromere) at the nuclear periphery. This can be achieved by yet unknown mechanisms of attraction between the NE proteins and specific histone modification and/or chromatin proteins. In this mechanism, the epigenetic profile of TADs determines both their affinity to the NE and transcriptional repression. That would explain both the presence of the negative correlation of transcription levels of L-TADs with the probability to be near the NE and the absence of any causal connection to transcription. In this picture, we assume that the state of being in affinity to the NE is set outside the interphase, at least outside the G1 state that we model. Second, the NE may contain proteins that interact with gene-poor chromatin and define the epigenetic status of TADs by modifying their histone tails. In this mechanism, the initial postmitotic interaction of a chromatin locus with the NE proteins sets its “LAD status” for the rest of the interphase. These settings include the accumulation of repressive epigenetic marks and frequent contact with the NE during the interphase. Experimental support exists for both of these mechanisms, see Discussion.

A way to reconcile our general findings with the tethering experiments described in the introduction is to assume that the epigenetic state of a TAD can be reset by contact with the lamina, but that reset requires a cell going through mitosis and forming a new NE, after which the locus has a relatively long and persistent contact with the NE. Likewise, once the epigenetic state is set, a long and persistent absence of contact is needed for a return to the original state. Only a subset of TADs is liable to such a reset. The underlying assumption is that a relatively long time is needed to reset the epigenetic state, as appears to be the case in experiments where the epigenetic state is reset by mechanical compression of the nucleus ***Wang et al.*** (***2019***).

## Materials and Methods

### Normalized measure of gene transcription levels at TAD resolution

RNA-seq methods generate data that needs to be normalized to eliminate technical biases associated with the methods, such as the sequencing depth of a sample and the length of the mRNA transcripts ***Wagner et al.*** (***2019***). To correct these biases, RPKM (reads per kilobase of transcript per million reads mapped) measure ***Mortazavi et al.*** (***2019***) has been widely used:

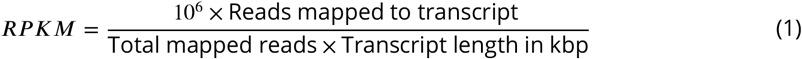

To quantify the average gene expression levels in a TAD, we propose a similar measure – RPKMT (number of reads mapped to all genes in a TAD per kilobase of the total length of all genes in a TAD per million reads mapped). This metric normalizes for the sum of all gene lengths in a TAD and the sequencing depth of the sample. RPKMT characterizes an average expression of all genes in a TAD, and is defined as:

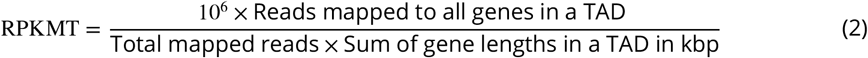

#### Calculation of RPKMT from published RNA-seq data

The RNA-seq data used here contains two replicates (rep1 and rep2) of control S2 cells (samples GSM3449348 and GSM3449349) or lamins depleted cells (samples GSM3449350 and GSM3449351) ***Ulianov et al.*** (***2019***). The TADs and their genomic coordinates are defined in ***Sexton et al.*** (***2019***). Annotated genes are assigned to a TAD if their genomic start positions are located within the TAD, according to BDGP Release 5.12/dm3. As a result, there are 8 out of 1169 TADs that do not contain any genes, therefore, in this work, we deliberately set the number of reads in these TADs to zero. Taking the two replicates into account, we calculate the transcription activity metric (RPKMT), Eq. 2, as:

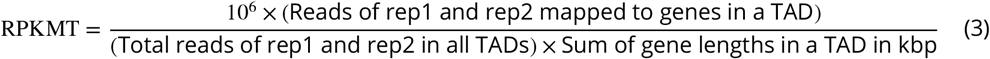

The FlyBase IDs of the genes (FBgn#), the genomic coordinates of the genes on chromosomes, and the genomic lengths of the genes (label “length”) are extracted from the fasta file: dmel-all-gene-r5.12.fasta (http://ftp.flybase.net/releases/FB2008_09/dmel_r5.12/fasta/).

We have verified that the proposed metric of transcription activity, Equation 3, used with the RNA-seq data from Ref. ***Ulianov et al.*** (***2019***), is consistent with the transcription activities in different epigenetic classes of TADs ***Sexton et al.*** (***2019***), which are employed by our computational model. The consistency is demonstrated in Figure 5 (see also Figure 5 in Appendix 1). Specifically, we consider two transcription profiles to be consistent if their corresponding medians over TAD classes satisfy the following criterion: the transcription level of Active TADs is much higher than those of all other epigenetic TAD classes. This consistency check mitigates potential concerns related to inevitable, but apparently minor, differences in gene expression profiles that may stem from differences between the types of embryonic cells from Ref. ***Ulianov et al.*** (***2019***) (S2 embryonic cell line) vs. those ***Sexton et al.*** (***2019***) (embryos collected 16-18 hour after egg laying) employed to build the dynamic computational model employed here. Another possible source of the observed differences in gene expression profiles shown in Figure 5, specifically a lower ratio of Active to non-Active median TAD levels in our RPKMT-based transcription profile (Figure 5A) compared to the profile from ***Sexton et al.*** (***2019***) (Figure 5B), may be related to the fact that, by construction, RPKMT metric normalizes for both the coding and non-coding regions of the genes that are located in a TAD. Moreover, due to the overlapping of the genomic regions of genes in a TAD and the way how the genes are assigned to a TAD (i.e., by their genomic start positions), the sum of the gene lengths in a TAD sometimes can exceed the length of a TAD.

### The dynamic model of interphase chromosomes at TAD resolution

To determine the probabilities of TADs to be in contact with the NE, we use a dynamic model of a *D*.*melanogaster* female interphase nucleus with a diploid set of four homologous chromosomes, developed in our previous work ***Tolokh et al.*** (***2019***). A brief description of the model is given below. The model simulates dynamics of the chromatin fibers in both control and lamins-depleted nuclei for a time equivalent to 11 hours (the duration of the *D*.*melanogaster* nucleus interphase) and allows analyzing the trajectories of individual TADs in both control and lamins-depleted nuclei. Langevin dynamics simulations have been performed using ESPResSo 3.3.1 package ***Limbach et al.*** (***2019***) by solving the Langevin equations of motion.

In the model, each pair of homologous chromosomes (2, 3, 4, and X), which are in proximity to each other ***Fung et al.*** (***2019***), is represented by a single chain of spherical beads using the bead-spring model ***Lifshitz et al.*** (***2019***); ***Mirny*** (***2019***). The four chains of these beads are surrounded by a spherical boundary representing the NE. 1169 beads in these chains correspond to 1169 pairs of homologous TADs - physical domains resolved in the Hi-C maps ***Sexton et al.*** (***2019***). Additionally, 4 beads represent centromeric chromatin domains (CEN) in each chain, and 6 beads adjacent to CEN beads represent pericentromeric constitutive heterochromatin domains (HET). The mass and the size of each bead correspond to the length of the DNA contained in the corresponding TAD, CEN, or HET domains.

The model employs four well-established major classes of TADs (Active, Null, PcG, and HP1) identified previously ***Sexton et al.*** (***2019***) based on their epigenetic signatures ***Filion et al.*** (***2019***) and biological functions. Following bead–TAD equivalence in the model, it has four corresponding bead types. Each bead/TAD type is characterized by its own interaction well depth parameter *ϵ*_*t*_ for attractive interactions between beads of the same type. Interactions between beads of different types are not type-specific and are characterized by a single well-depth parameter *ϵ*_*g*_. The beads corresponding to TADs that contain lamina-associated domains (LADs) ***Pickersgill et al.*** (***2019***); ***van Bemmel et al.*** (***2019***) – L-TADs – can attractively interact with the NE. This L-TAD–NE affinity can temporally confine L-TADs at the NE. The interaction parameters of the model are tuned to reproduce: (i) the average experimental fraction of LADs confined to the NE, 25% ***Pickersgill et al.*** (***2019***) and (ii) the experimental TAD-TAD contact probability (Hi-C) map ***Sexton et al.*** (***2019***); ***Li et al.*** (***2019***) (Pearson’s correlation coefficient is 0.956). The model predicts highly dynamical distributions of the chromatin (both for control and lamins-depleted cells), which, after averaging, are in good agreement with the experimentally observed average density profiles of fruit fly chromatin ***Bondarenko and Sharakhov*** (***2019***). As in the experiment, the chromatin density distribution in the model lamins-depleted cells shows a substantial shift of the chromatin from the NE, compared to the distribution in control nuclei, accompanied by a large increase of the density in the central nucleus region (see Figure 2 of Appendix 1). The model also correctly reproduces the experimentally observed changes of average radial positioning of individual cytological regions (22A, 36C, and 60D) explored previously ***Ulianov et al.*** (***2019***) in the lamins-depleted cells, Figure 1 of Appendix 1. The model is validated against multiple features of chromatin structure from several experiments not used in model development.

The probability of a TAD being in contact with the NE is calculated as the probability of the center of the corresponding bead being within 0.2 *μ*m spherical layer adjacent to the NE. The thickness of this layer is barely larger than the radius of the largest TAD (0.19 *μ*m) and corresponds to the average TAD diameter in the nucleus model used. To calculate the probabilities, we analyzed 6 trajectories of model nuclei (corresponding to different nucleus chromatin topologies), each containing 400 × 10^3^ snapshots (chromatin configurations).

### Binning of TADs based on their probabilities to be in contact with the NE

The TADs are grouped into six bins, according to the probability of each TAD in control cells to be in contact with the NE. For 1169 TADs, Figure 3A and D, bins #1–5 contain 194 TADs each, whereas bin #6 has 195 TADs. For 350 L-TADs, Figure 3C and F, bins #1-5 contain 58 L-TADs each, whereas bin #6 has 60 L-TADs. For 819 NonL-TADs, Figure 3B and E, bins #1–5 contain 137 nonL-TADs each, whereas bin #6 has 134 nonL-TADs. The number of bins is determined by a reasonable balance between two opposing requirements: enough bins are needed to clearly identify a trend in transcription level changes, but too many bins result in a higher standard error of the mean transcription level per bin.

## Acknowledgments

This research was supported by the National Science Foundation (MCB-1715207). A.V.O. acknowledges partial support from the NIH GM144596. A.V.O. thanks Andrei Onufriev for implementing and running the numerical example supporting the gene activity ratio analysis.

## Availability of software

The modeling code ESPResSo 3.3.1 ***Limbach et al.*** (***2019***) used in this research is available at http://espressomd.org/. The software is free, open-source, and published under the GNU General Public License (GPL3).

## Consent for publication

All authors read and approved the manuscript.

### Appendix 1

#### Distributions of TADs in the Drosophila nucleus

**Appendix 1 Figure 1.**
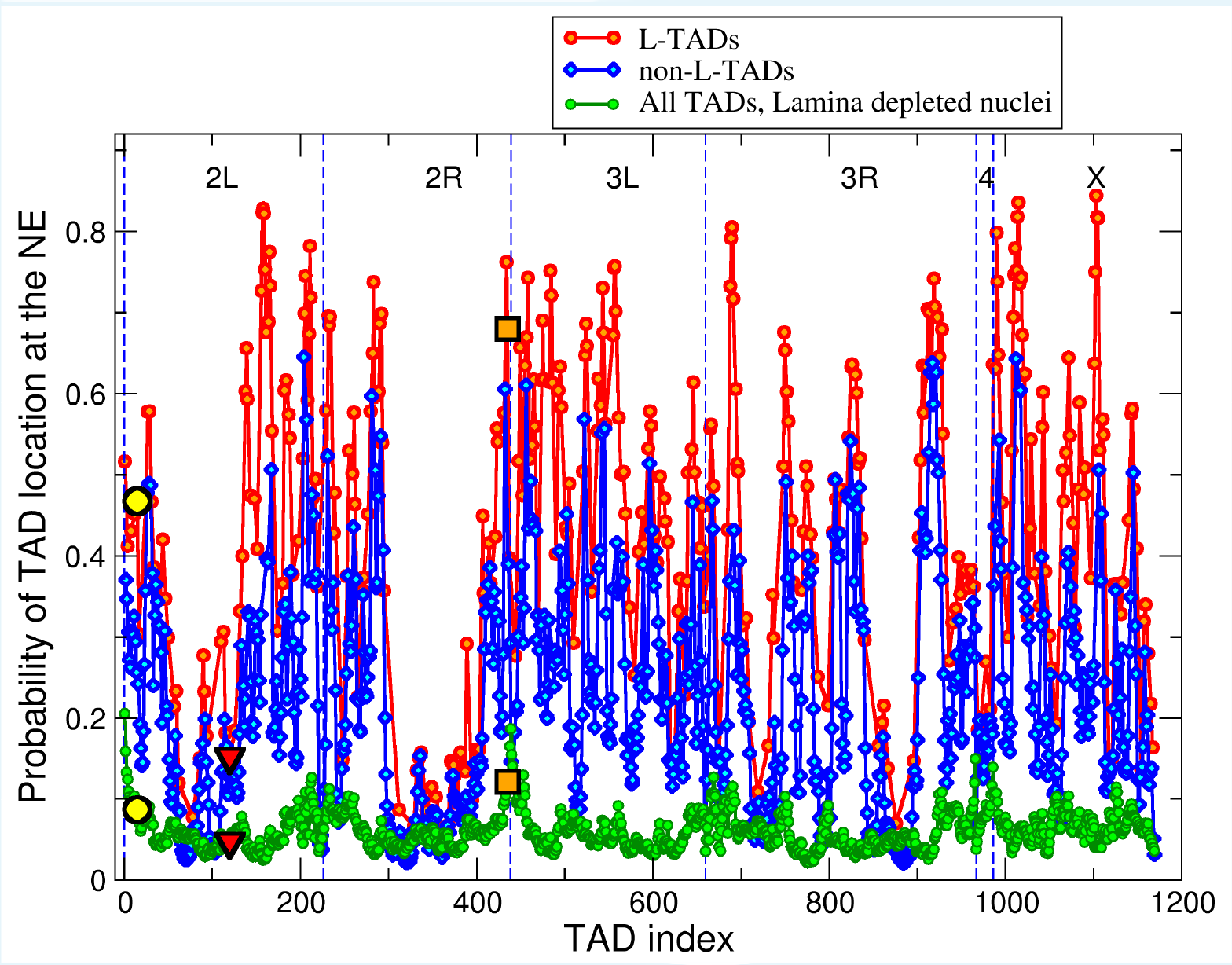
Probabilities of TADs (LAD containing TADs (L-TADs) and TADs not containing LADs (nonL-TADs) in control nucleus model, and all TADs in lamins depleted nucleus model) to be in contact with the NE (to be within 0.2 *μ*m from the NE). Null L-TAD #15 (in control and lamins depleted nuclei), analyzed in ***Ulianov et al.*** (***2019***) as cytological region 22*A*, is marked by yellow circles. Null L-TAD #120 (in control and lamins depleted nuclei), analyzed in ***Ulianov et al.*** (***2019***) as cytological region 36*C*, is marked by red triangles. PcG L-TAD #435 (in control and lamins depleted nuclei), analyzed in ***Ulianov et al.*** (***2019***) as cytological region 60*D*, is marked by orange squares.

**Appendix 1 Figure 2.**
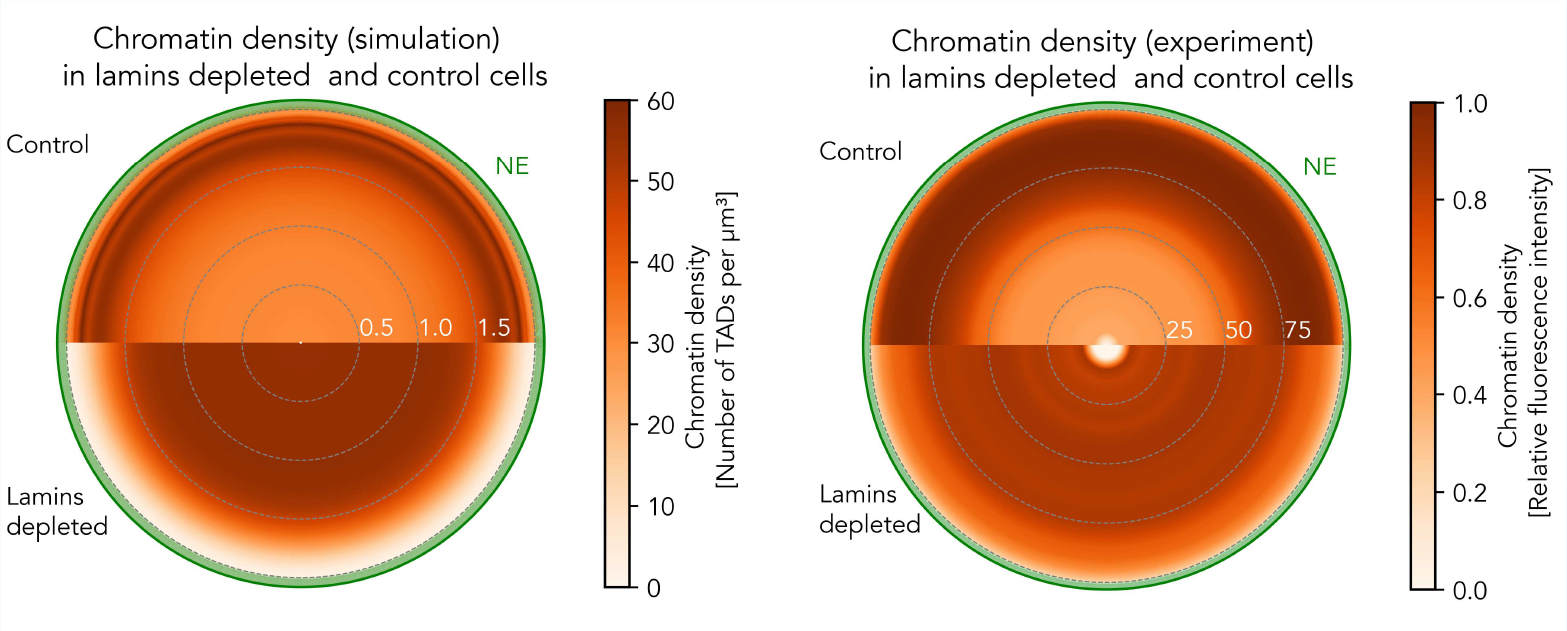
**Left panel**: Computed chromatin density averaged over the spherical layers as a function of the radial distance from the nucleus center in control nuclei (top) and in lamins-depleted nuclei (bottom). The radius of the nucleus is 2 *μ*m. **Right panel**: Experimental mean chromatin radial density in the equatorial plane of the nucleus of the proventriculus. For illustration only, the azimuthal dependence of the density is averaged out to produce a schematic that shows only the radial density profile. The density is inferred from relative fluorescence intensity, as detailed in Ref. ***Bondarenko and Sharakhov*** (***2019***). Specifically, 21 equally spaced experimental data points are taken from Fig. S3 (Group 1, bottom panel) of Ref. ***Bondarenko and Sharakhov*** (***2019***) and then interpolated using a linear interpolation process, yielding 201 equally spaced data points plotted in the figure. The radial position of the mean chromatin density is measured from the nuclear center to the periphery (0% - 100%).

#### Comparing averages and ratios of gene expression levels

We argue here that, counter-intuitively, the use of ratios of gene expression levels to characterize possible differences in transcription activities between two sets of genes (e.g. knockdown vs. control) can lead to unintended biases due to inherent noise in the data. For the sake of argument, consider a simplified case of two sets *C* and *K*, of *N* genes in each set, each gene having the same inherent transcription level in both sets. Due to the inevitable stochasticity of gene expression, especially relevant at low levels, and because of experimental uncertainty, the actual measurement of each gene activity will be a random variable 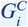 (and 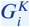) with some distribution, here assumed identical for all genes. For the sake of argument, assume this distribution to be uniform on the gene activity interval from 0 to 1. Obviously, in this case the mean expression level ⟨*G*_*i*_⟩ of each gene is exactly 1/2, the activity averaged over each gene set 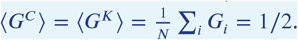. That is if one uses transcription activity averages to compare two sets of genes, their activities are the same, as expected. The situation is different if one attempts to use 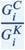 to make the comparison, *e*.*g*. to evaluate the effect of a knockdown. Note that, in general, the average of a ratio does not equal the ratio of the averages; a numerical example is shown in Table 1. The intuitive rationale for the effect is as follows: a deviation of the denominator down from its mean value causes a larger increase of the fraction than does the decrease of the fraction caused by the same size deviation of the denominator up from its mean. Rigorous analysis shows that in the case of the uniform distribution on [0 1] interval, the mean of the ratio diverges (logarithmically), which explains the large variation of the mean ratio from one trial (“experiment”) to another. Thus, each independent set of measurements can bring about a different outcome in terms of the ratio of the gene activities, Table 1.

**Appendix 1 Table 1.**
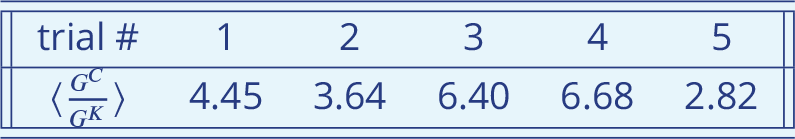
A numerical simulation of gene activity with noise. Here, *G*^*C*^ and *G*^*K*^ are uniformly distributed random variables on the interval [0,1]. A total of 2*N* = 2000 random numbers were generated for each trial, and ratios of two sequential random numbers were computed and averaged over all *N* pairs. Each trial starts with an independent seed to initiate the random number generator `Math.random()`, as implemented in Java 1.16.4.

| trial # | 1 | 2 | 3 | 4 | 5 |
| --- | --- | --- | --- | --- | --- |
| $\langle \frac{G^C}{G^K} \rangle$ | 4.45 | 3.64 | 6.40 | 6.68 | 2.82 |

#### The key conclusions remain valid with respect to another metric of transcription activity in TADs

We propose another metric of transcription activity in TADs -RPKMTL (number of Reads mapped to all genes in a TAD Per Kilobase of TAD length per Million reads mapped to all TADs). Unlike RPKMT, RPKMTL uses the length of a TAD to obtain the average transcription activity at TAD resolution. RPKMTL characterizes an average expression of all genes in a TAD and is defined as:

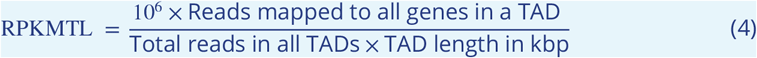

For two replicates (rep1 and rep2) from published RNA-seq data ***Ulianov et al.*** (***2019***), the transcription activity metric, Equation 4, is calculated as:

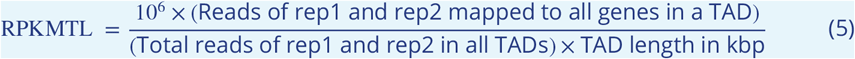

**Appendix 1 Figure 3.**
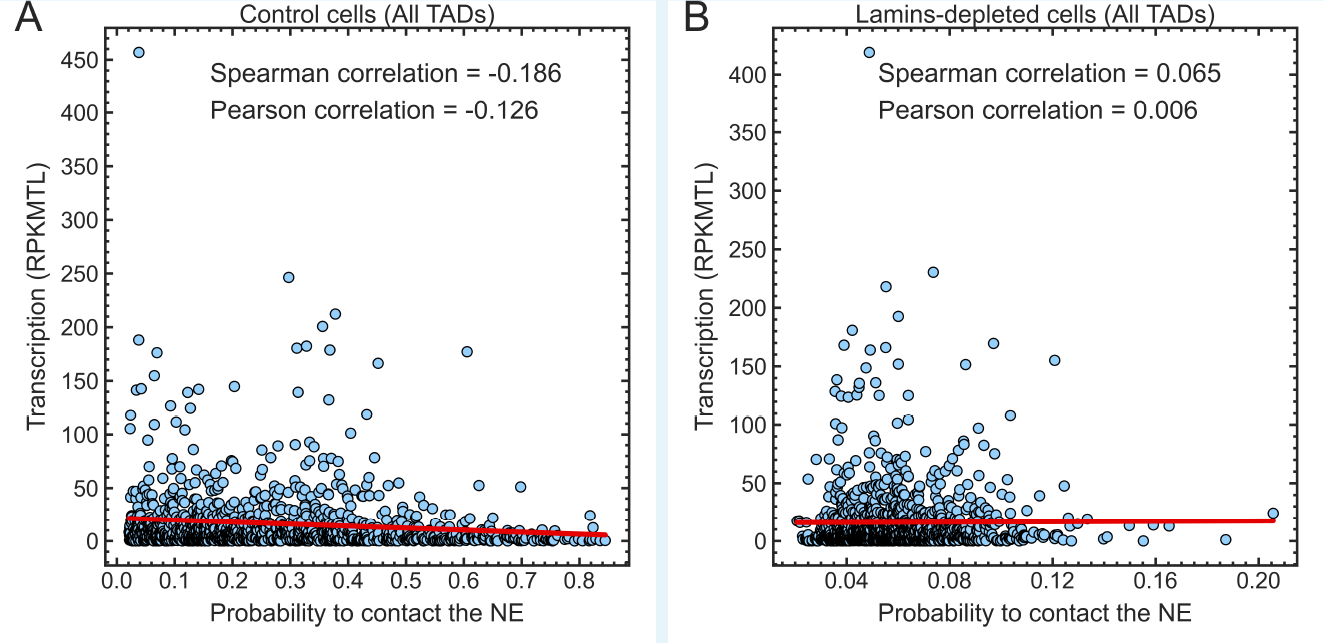
**(A)** Scatter plot shows a weak negative correlation between the expression of genes in TADs (in RPKMTL) and the probability of TAD to be found in contact with the NE (i.e. to be found within 0.2 *μ*m layer near the NE) in the control nuclei. **(B)** Scatter plot shows essentially no correlation between the TAD expression (in RPKMTL) and the probability of TAD being found in contact with the NE in the lamins-depleted nuclei. The Spearman, Pearson correlation coefficients and linear regression lines (red) are shown. Plots are generated with the *lmplot* function in Seaborn.

**Appendix 1 Figure 4.**
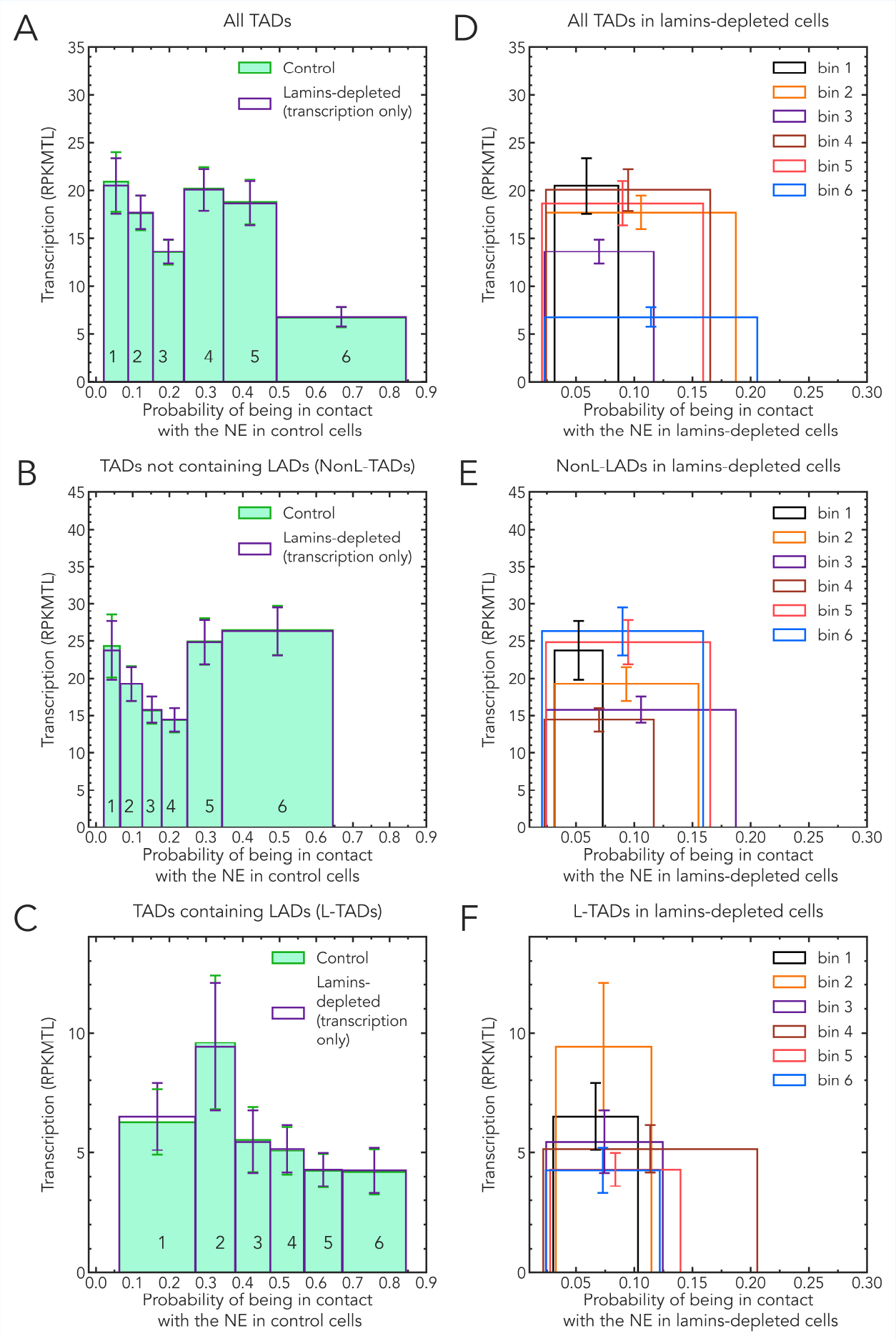
Dependencies of bin averaged TAD transcription levels (RPKMTL) on the average probability of the TADs in the bin to be in contact with the NE. The binning of TADs is based on TAD-NE contact probabilities in control cells for each set (selection) of TADs. **Solid bars: control cells. Empty bars: lamins depleted cells**. The same set of TADs per bin is used in the control and lamins-depleted cells. Error bars are s.e.m. (standard error of the mean). **Left panels: (A)** all TADs; **(B)** TADs not containing LADs (NonL-TADs); and **(C)** TADs containing LADs (L-TADs). In the left panels only, the positions of the empty bins (lamins depleted cells) along the x-axis are deliberately kept unchanged to facilitate visual comparison with the heights of the corresponding control bins. **Right panels: (D)** all TADs; **(E)** TADs not containing LADs (NonL-TADs); and **(F)** TADs containing LADs (L-TADs). A clear shift of the average TAD positions away from the NE is evident.

**Appendix 1 Figure 5.**
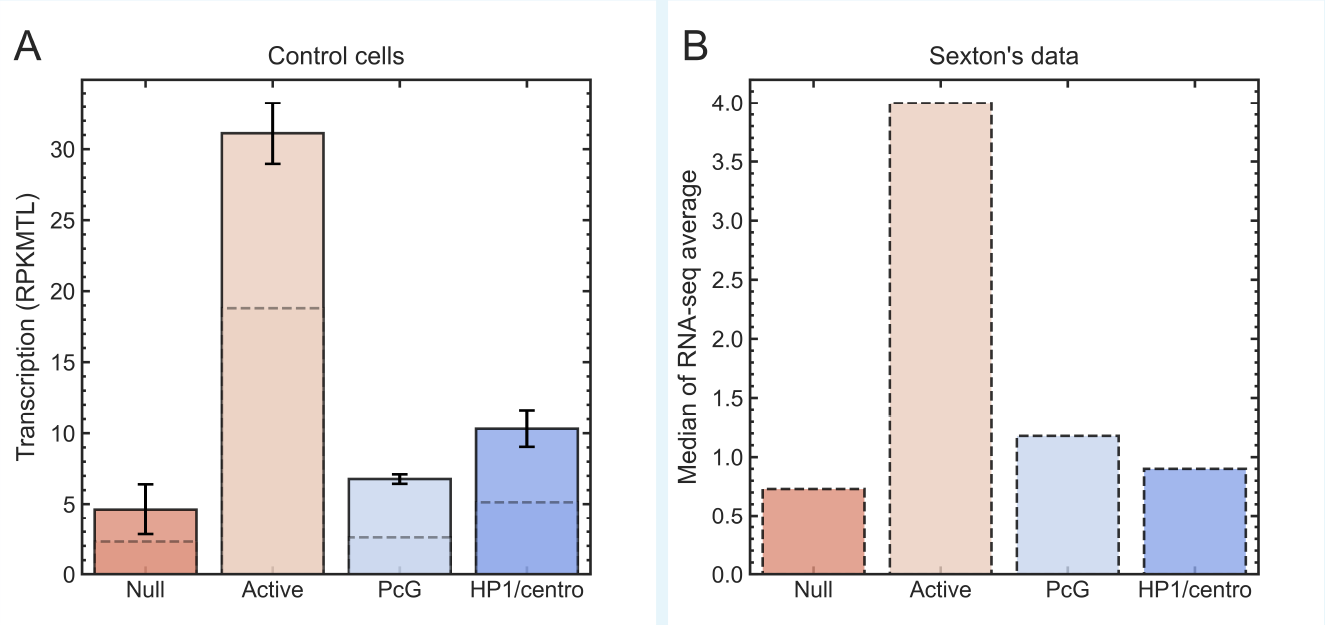
The metric of transcription activity in TADs, RPKMTL, is consistent with the epigenetic classes of TADs identified previously ***Sexton et al.*** (***2019***). Median transcription level (in RPKMTL) in Active TADs (n=494) is at least 2 times greater than those of other epigenetic TAD classes, such as HP1/centromeric (n=52), Null (n=492), and PcG (n=131) (**panel A, dashed lines**). The medians (dashed lines) along with the means (solid boxes) demonstrate consistency with the data in Figure 3C of Ref. ***Sexton et al.*** (***2019***), reproduced in the **panel B**, which show the median gene transcription levels within each epigenetic class of TADs. The dynamic model of fruit fly nucleus employs the partitioning of genome into TADs and their epigenetic classes, introduced in Ref. ***Sexton et al.*** (***2019***). Error bars are s.e.m. (standard error of the mean).

#### Distribution of TADs by types and epigenetic classes

**Appendix 1 Figure 6.**
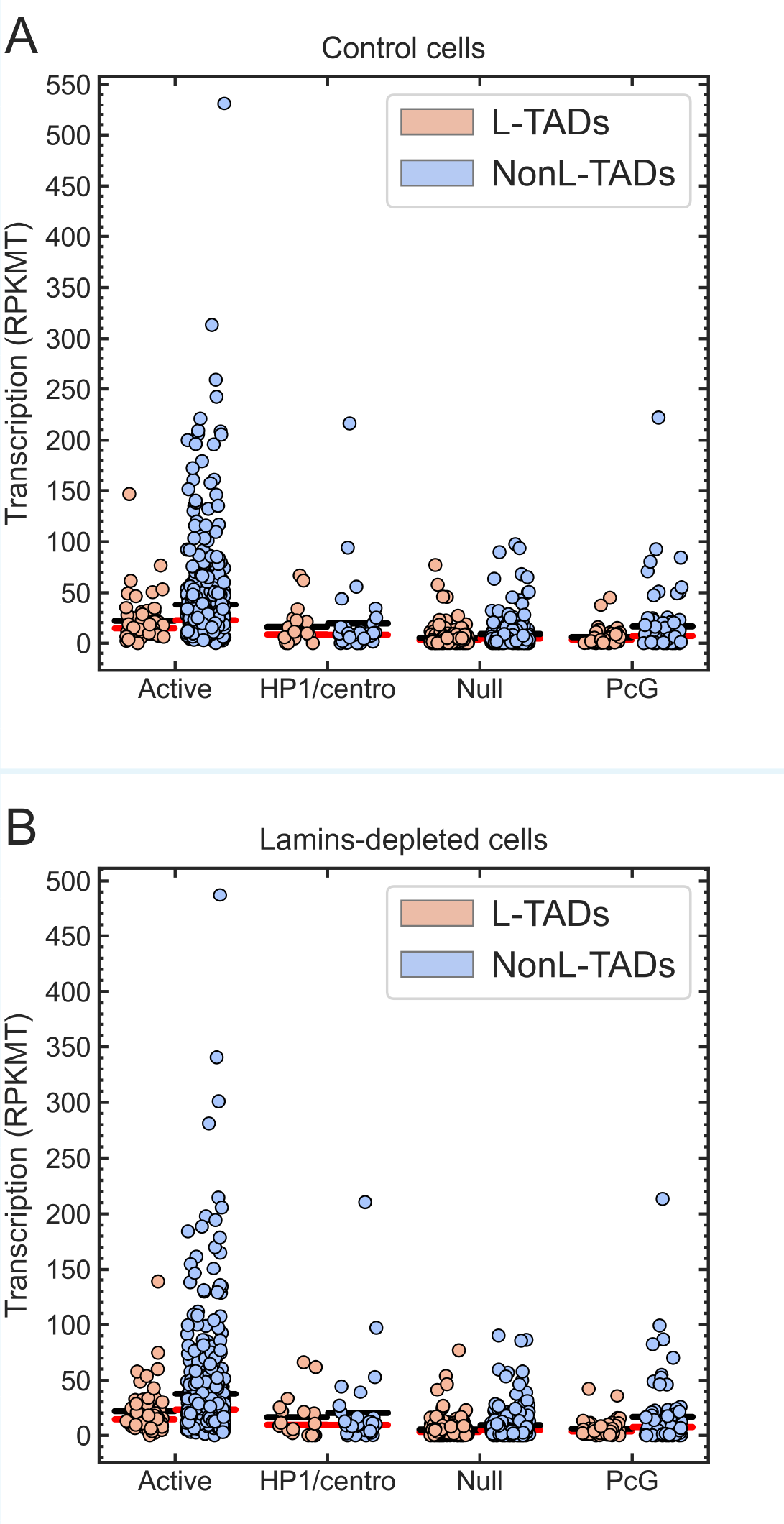
Distribution of TADs by types such as Active L-TADs (n=54), Active NonL-TADs (n=440), HP1/centromeric L-TADs (n=34), Null L-TADs (n=228), Null NonL-TADs (n=264), PcG L-TADs (n=50), and PcG NonL-TADs (n=81) in control (**A**) and lamins-depleted (**B**) cells. The average gene expression (black horizontal lines) in L-TADs for each type is lower than those of NonL-TADs. The red horizontal lines are the median gene expression values in TADs for each type.

#### A note on bins #3 and #4 in Figure 3B

Below is a possible explanation for why the average transcription levels (in RPKMT) of NonL-TADs in bins #3 and #4 in Figure 3B, corresponding to the 0.15–0.28 probabilities of NonL-TADs to be in contact with the NE, are relatively low. Here we compared the fractions of different epigenetic classes of TADs in each of the six bins in Figure 3B. Bins #3 and #4 demonstrate a reduced number of Active TADs, which have a much higher average transcription level compared to other three (non-Active) epigenetic classes of TADs (see Figure 5), and increased fractions of non-Active TADs, see Figure7E of Appendix 1 below. In contrast, bin #5 and bin #6 (relatively high average RPKMT levels in Figure 3B) have higher fractions of Active TADs relative to other bins (see Figure7E of Appendix 1).

**Appendix 1 Figure 7.**
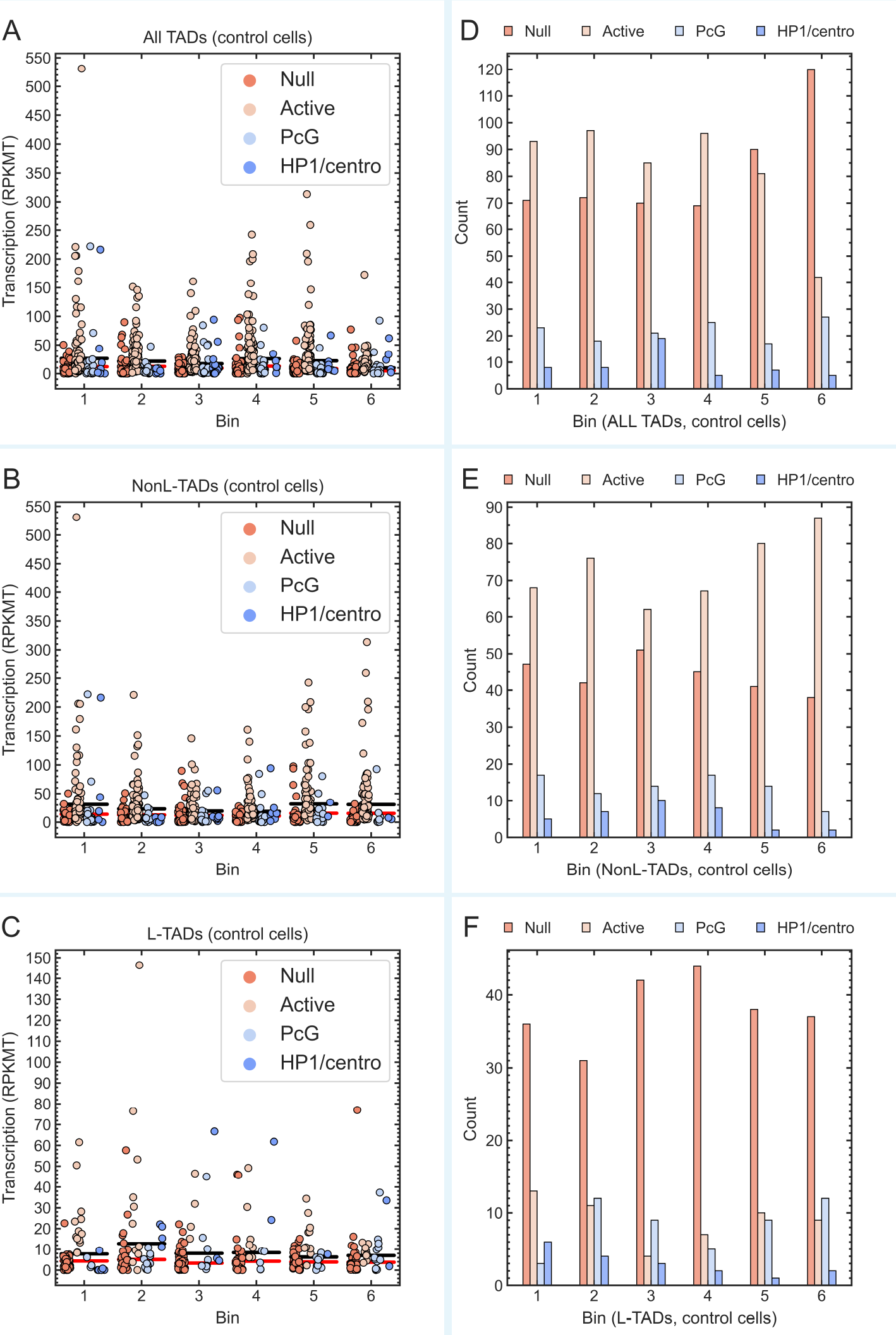
Distribution of TADs in bins by epigenetic classes (Null, Active, PcG, and HP1/centromeric). **(A, B, and C)** The red and black horizontal lines are the median and mean gene expression values (in RPKMT) in each bin, respectively. **(D, E, and F)** Comparison of the number of TADs of each epigenetic class in each bin. For NonL-TADs (control cells), the number of Active TADs is greater than those of other epigenetic classes in each bin. In contrast, for L-TADs (control cells), the number of Null TADs is greater than those of other epigenetic class in each bin. Positions of TADs along the horizontal axis in bins are not to scale.

